# MQF and buffered MQF: Quotient filters for efficient storage of k-mers with their counts and metadata

**DOI:** 10.1101/2020.08.23.263061

**Authors:** Moustafa Shokrof, C. Titus Brown, Tamer A. Mansour

## Abstract

**Background:** Specialized data structures are required for online algorithms to efficiently handle large sequencing datasets. The counting quotient filter (CQF), a compact hashtable, can efficiently store k-mers with a skewed distribution.

**Result:** Here, we present the mixed-counters quotient filter (MQF) as a new variant of the CQF with novel counting and labeling systems. The new counting system adapts to a wider range of data distributions for increased space efficiency and is faster than the CQF for insertions and queries in most of the tested scenarios. A buffered version of the MQF can offload storage to disk, trading speed of insertions and queries for a significant memory reduction. The labeling system provides a flexible framework for assigning labels to member items while maintaining good data locality and a concise memory representation. These labels serve as a minimal perfect hash function but are ~10 fold faster than BBhash, with no need to re-analyze the original data for further insertions or deletions.

**Conclusion:** The MQF is a flexible and efficient data structure that extends our ability to work with high throughput sequencing data.

## Background

Online algorithms effectively support streaming analysis of large data sets, which is important for analyzing data sets with large volume and high velocity(1). Approximate data structures are commonly used in online algorithms to provide better average space and time efficiency (2). For example, the Bloom filter supports approximate set membership queries with a predefined false positive rate (FPR) (3). The count-min sketch (CMS) is similar to Bloom filters and can be used to count items with a tunable rate of overestimation. However, there are a number of problems with Bloom filters and the CMS - in particular, they do not support data locality.

The Counting Quotient Filter (CQF) is a more efficient data structure that serves similar purposes with better efficiency for skewed distributions and much better data locality(4). The CQF is a recent variant of quotient filters that tracks the count of its items using a variable size counter. As a compact hashtable, CQF can perform in either probabilistic or exact modes and supports deletes, merges, and resizing.

Analysis of k-mers in biological sequencing data sets is an ongoing challenge(5). K-mers in raw sequencing data often have a high Zipfian distribution, and the CQF was built to minimize memory requirements for counting such items. However, this advantage deteriorates in applications that require frequent random access to the data structure, and where the k-mer count distribution may change in response to different sampling approaches, library preparation and/or sequencing technologies. For example, k-mer frequency across 1000s of RNAseq experiments shows different patterns of abundant k-mers (6).

Data structures like CMS (7) and CQF (4) also do not natively support associating k-mers with multiple values, which can be useful for coloring in De Bruijn graphs as well as other features (8). Classical hash tables are designed to associate their keys with a generic data type but they are expensive memory-wise (9). Minimal Perfect Hash Functions (MPHFs) can provide a more compact solution by mapping each k-mer into a unique integer. These integers can then be used as indices for the k-mers to label them in other data structures (10). An implementation capable of handling large scale datasets with fast performance requires ~3 bits per element (11). However, such a concise representation comes with a high false-positive rate on queries for non-existent items. Moreover, unlike hashtables, MPHF does not support insertions or deletions thus any change in the k-mer set would require rehashing of the original dataset.

In this paper, we introduce the mixed-counters quotient filter (MQF), a modified version of the CQF with a new encoding scheme and labeling system supporting high data locality. We further show how Buffered MQF can be used to scale MQF to solid-state disks. We compare between MQF and the CQF, CMS, and MPHF data structures regarding memory efficiency, speed performance, and applicability to specific data analysis challenges. We further do a direct comparison of the CMS to MQF in the khmer software package for sequencing data analysis, to showcase the benefits of MQF is in real world applications.

## Results

### MQF has a lower load factor than CQF

The load factor is defined as the actual space utilized divided by the total space assigned for the data structure, and is an important measure of data structure performance. To compare load factors between the CQF and MQF data structures, instances of both structures were created using the same number of slots (2^27^). Chunks of items from five datasets with different distributions of item frequencies were inserted iteratively to both data-structures while recording the load factor after the insertion of each chunk. The experiments stopped when MQF’s load factor reached 90%. MQF had lower loading factors for all tested datasets but the difference was minimal for the dataset with the highest Zipfian distribution (Z=5). The lower the tested Zipfian distribution the lower the loading factor of MQF (Figure 1). A lower loading factor enabled MQF to accommodate > 30% of the CQF capacity from a dataset of real k-mers and to exceed the double CQF capacity with uniform distribution (Figure 1 and supplementary table 1).

**Figure 1:**
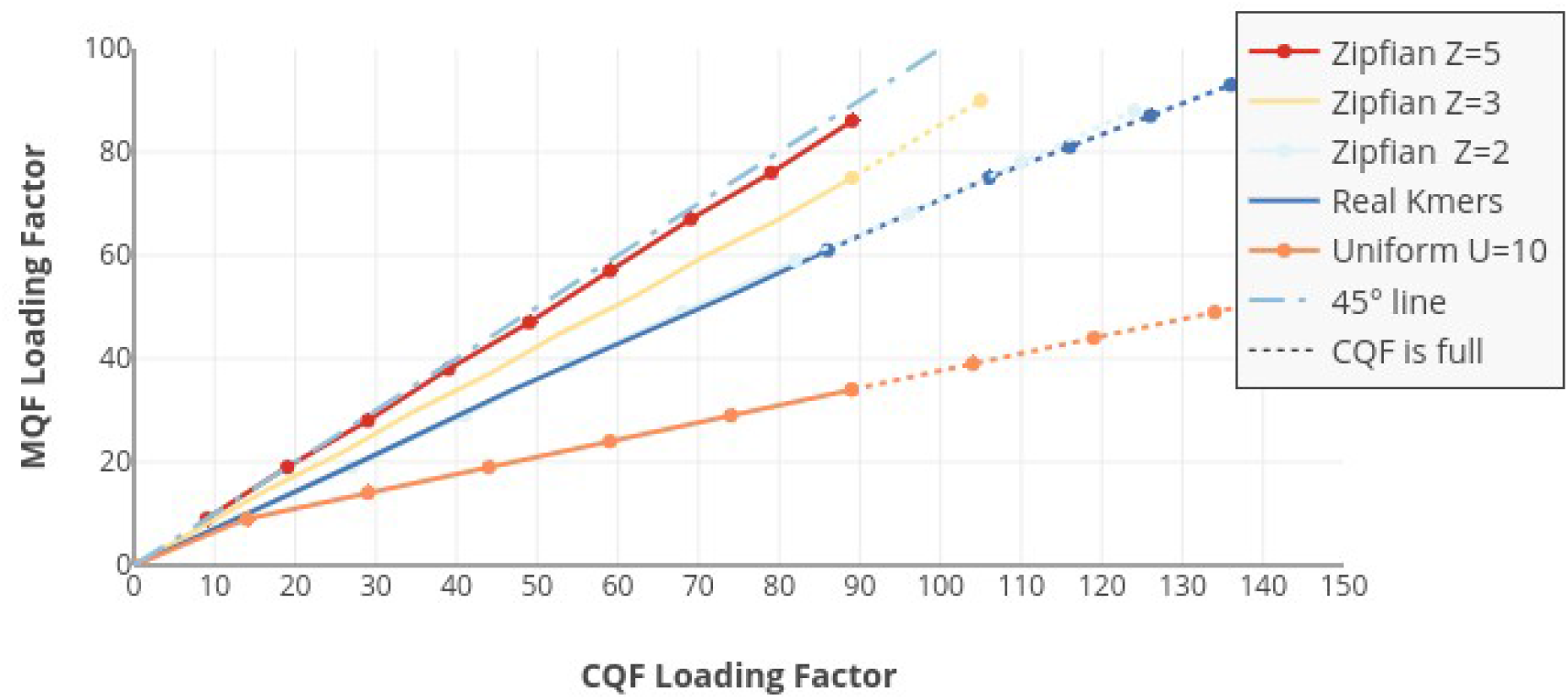
MQF has a lower load factor compared to CQF. Chunks of items, from different distributions of item’s frequencies, were inserted iteratively to matching CQF and MQF structures. MQF had lower loading factors for all tested datasets with better performance with more uniform distributions (The further from the 45° line the better the MQF).

### MQF is usually more memory efficient than CQF

Progressively increasing numbers of items were sampled from the real and Zipfian-simulated datasets. The smallest CQF and MQF to store the same number of items from each dataset were created. To do that, the q parameter of CQF versus the q and F_size_ parameters of MQF were calculated empirically. MQF was more memory efficient for real k-mers and Zipfian-simulated distributions with low coefficients in 75% of the cases (Figure 2). The tuning of the F_size_ enabled MQF to grow in size gradually compared to CQF which has to double in size to fit the minimal increase in items beyond the capacity of a given q value (Supplementary Figure 1).

**Figure 2:**
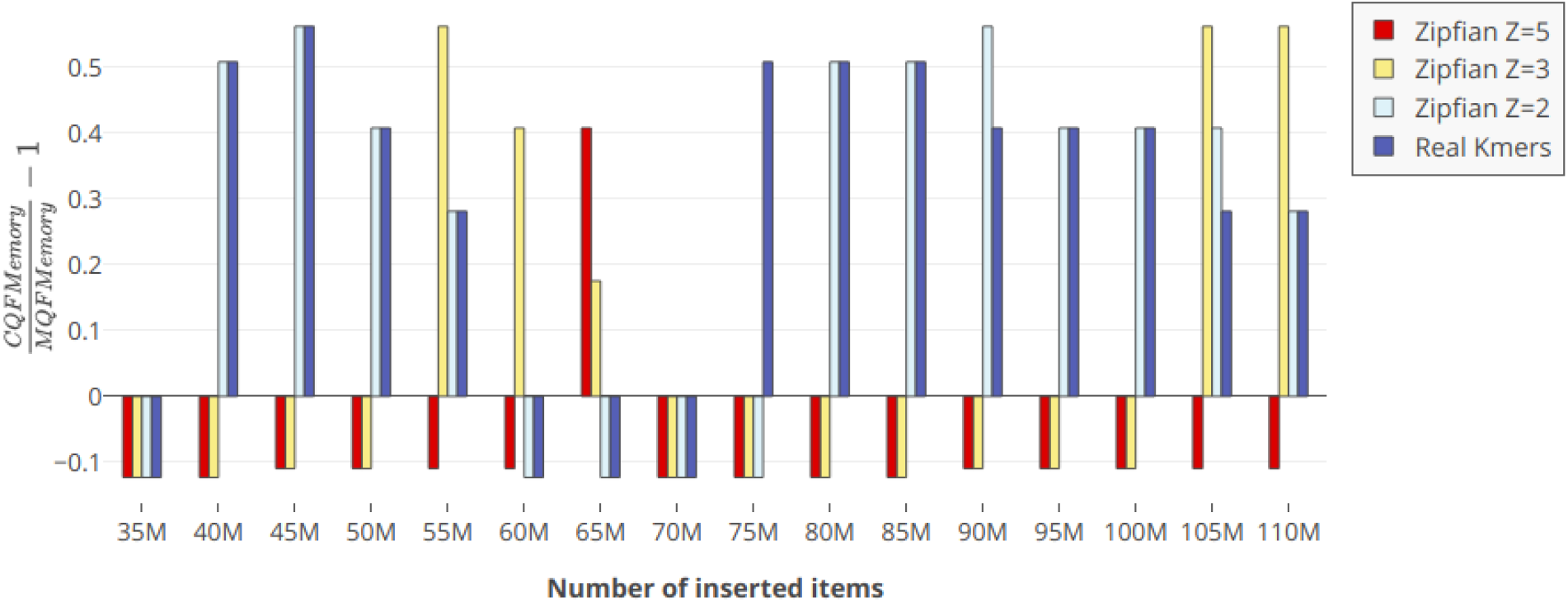
Memory consumption comparison between CQF and MQF. The graph compares the memory consumption of the smallest CQF and MQF that fits different datasets. The bigger the value on the y-axes, the more memory the MQF saved.

### MQF is faster than CQF and low-FPR CMS

The in-memory and buffered MQFs were evaluated for speed of insertion and query in comparison to three in-memory counting structures: CQF, the original CMS (12), and khmer’s CMS (13). To test the effect of FPR on the performance, the experiment was repeated for 4 different FPRs (0.1, 0.01, 0.001, 0.0001). All tested structures were constructed to have approximately the same memory space except for buffered MQF which used only one-third of this memory for buffering while the full-size filter is on the disk. MQF is guaranteed to hold the same number of items as a CQF having the same number of slots. The number of slots in CQF was chosen so that the load factor was more than 85% and the MQFs were created with an equal number of slots. Items were sampled for insertion from the real and Zipfian-simulated datasets. After finishing the insertion, to assess the query rate, 5M items from the same distribution as the insertion datasets were queried. Half of the query items didn’t exist in the insertion datasets.

MQF has a faster insertion and query rates compared to CQF with minimal, if any, effect of the FPR on either structure. The performance of CMS is better with higher FPR and Khmer’s implementation of CMS doubles the query rate of the original one. However, MQF is always faster than both CMS unless the FPR is more than 0.01 (Figure 3).

**Figure 3:**
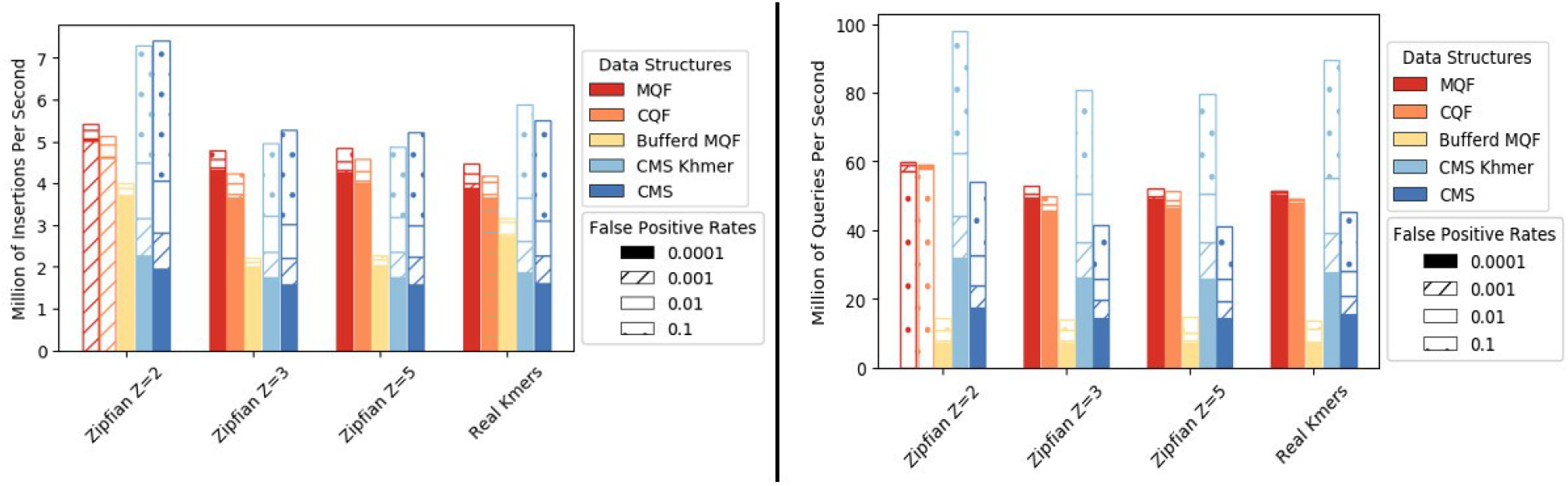
Performance comparison of four data-structures: MQF, CQF, buffered MQF, Khmer implementation of CMS, and Original implementation of CMS: Insertions rate (left panel) and query rate (right panel).

### MQF outperforms CMS in real-world problems

Khmer is a software package deploying a new implementation of CMS for k-mer counting, error trimming and digital normalization (13). To test MQF in real-life applications, we assessed the performance of the Khmer software package using CMS (13) versus our new implementation using MQF (https://github.com/dib-lab/khmer/tree/MQFIntegration2). A real RNA seq dataset with 51 million reads from the Genome in a Bottle project (14) was used for error trimming and digital normalization; two real-world applications that involve both k-mer insertions and queries. An exact MQF was used to create a benchmark for the approximate data structures. It took 5Gb RAM to create the data structure and 45 and 43 minutes to perform trimming and digital normalization respectively. The optimal memory for MQF and the optimal number of hash functions for CMS were calculated to achieve the specified false-positive rates. The CMS was constructed with the same size as the corresponding MQFs. The CMS and MQF versions of Khmer were compared regarding the speed and accuracy (Table 1).

**Table 1:** Khmer performance in error trimming and digital normalization using MQF and CMS. *Percentages of wrong decisions made by CMS at FPR = 0.1 in error trimming and digital normalization are 0.8% and 0.13% of the total number of decisions versus 0.02% and 0.01% made by MQF.

| FPR | Memory in GB | Error Trimming |  |  |  | Digital normalization |  |  |  | Error Bound in CMS | Hash func. In CMS |
| --- | --- | --- | --- | --- | --- | --- | --- | --- | --- | --- | --- |
|  |  | Time in Min. |  | Missed reads with Errors |  | Time in Min. |  | Reads kept by Error |  |  |  |
|  |  | MQF | CMS | MQF | CMS | MQF | CMS | MQF | CMS |  |  |
| 10<sup>-1</sup> | 1.8 | 42 | 39 | 11011 | 445817* | 39 | 37 | 3253 | 31143* | 13,11 | 3 |
| 10<sup>-2</sup> | 2.6 | 43 | 48 | 1304 | 404354 | 41 | 45 | 416 | 24987 | 14 | 5 |
| 10<sup>-3</sup> | 3.4 | 44 | 61 | 130 | 311464 | 42 | 54 | 58 | 21000 | 15 | 7 |
| 10<sup>-4</sup> | 4.5 | 44 | 75 | 3 | 292746 | 42 | 68 | 4 | 18449 | 16 | 10 |
| Exact | 5 | 45 | - | 0 | - | 43 | - | 0 | - | - | - |

### MQF is faster than MPHF

MPHF is constructed by default to fit the input k-mers while MQF would have different load factor that might affect its performance. To address this question, four growing subsets of real k-mers were inserted into MQFs of size 255 MB to achieve 60%, 70%, 80%, and 90% load factors. MPHFs were constructed with sizes ranging from 15 to 22 MB to fit the four datasets. All data structures were queried with 35M existing k-mers and the query times were reported. The MQFs were ~10 folds faster than the MPHFs. The query time of the MQF was invariable over the different load factors (Supplementary Figure 2).

## Discussion

MQF is a new variant of counting quotient filters with novel counting and labeling systems. The new counting system increases memory efficiency as well as the speed of insertions and queries for a wide range of data distributions. The labeling system provides a flexible framework for labeling the member items while maintaining good data locality and a concise memory representation.

MQF is built on the foundation of CQF. MQF has the same ability to behave as an exact or approximate membership query data structure while tracking the count of its members. The insertion/query algorithm developed for CQF enables this family of compact hashtables to perform fast under high load factor (up to 95%) (4). CQFs are designed to work best for data from high Zipfian distributions. However, previous k-mer spectral analysis of RNAseq datasets showed substantial deviations from a Zipfian distribution in thousands of samples(6). Such variations in distribution are expected given the variety of biosamples, the broad spectrum of sequencing techniques, and different approaches to data preprocessing.

MQF implements a new counting system that allows the data structure to work efficiently with a broader range of data distributions. The counting system adopts a simple encoding scheme that uses a fixed small space alone or with a variable number of the filter’s slots to record the count of member items (Figure 4). Items with small counts utilize the small fixed-size counters. Therefore, slots, used to be consumed by CQF as counters for these items, are freed to accommodate more items in the filter. The MQF’s load factor grows slower than CQF with all distributions except the extreme Zipfian case (Z=5) where the load factor is almost the same (Figure 1). This is why the memory requirement for MQFs is usually smaller compared to CQFs under most distributions despite the extra space taken by the fixed counters (Figure 2). The size of the fixed-size counter is constant independent from the slot size, therefore the memory requirement for this counter will be trivial with big slots for smaller FPRs and almost negligible in the exact mode. However, this fixed-size counter comes with an additional advantage for MQF. Tuning the size of the fixed-size counter enables the filter to accommodate more items with a slightly larger slot size. This allows the memory requirement for MQF to grow gradually instead of the obligatory size doubling seen in CQF (Figure 2 and Supplementary Figure 1).

**Figure 4:**
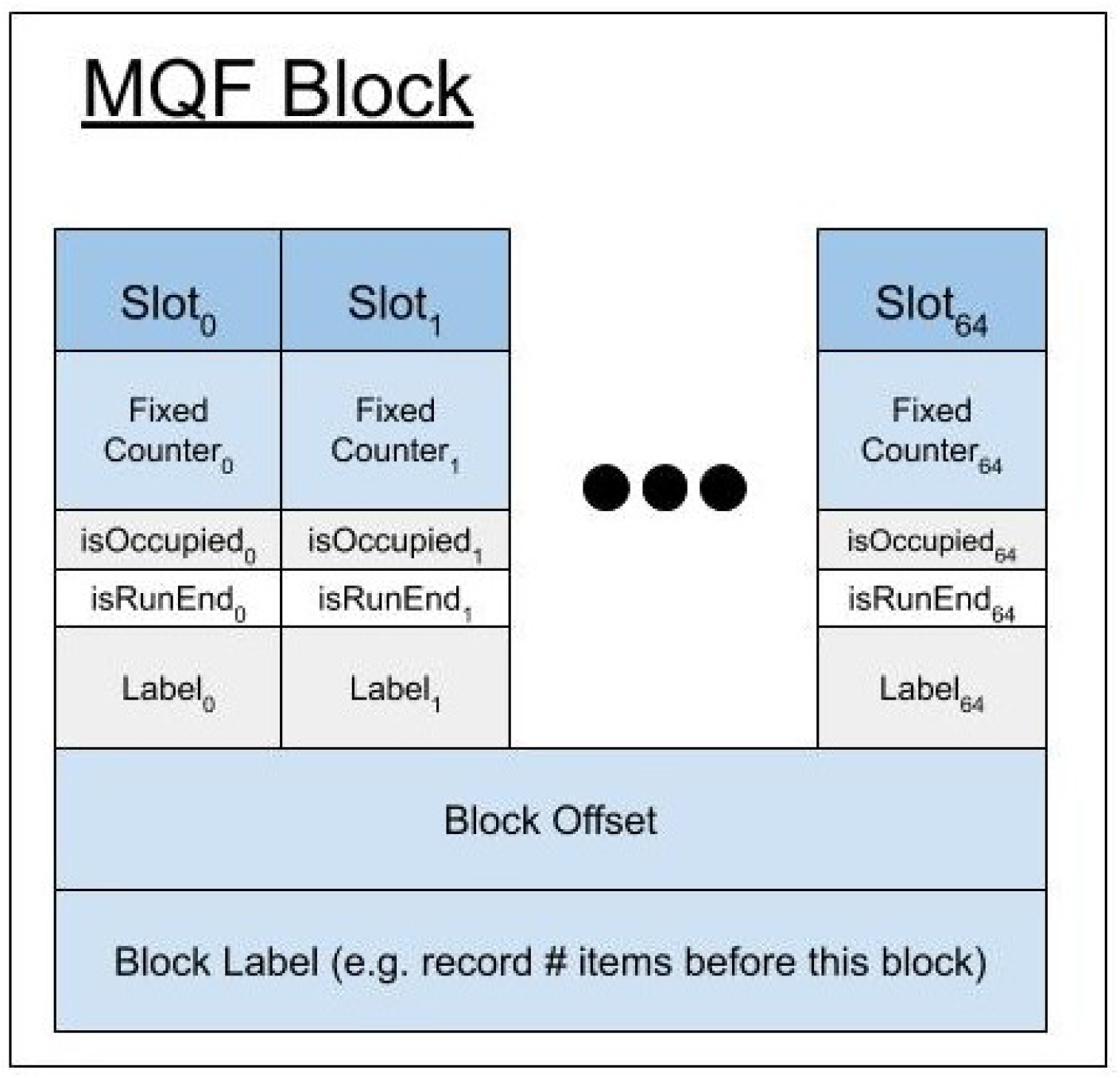
MQF block structure. Each MQF block contains 64 slots with their metadata, a one-byte block offset, and configurable size space to hold the number of items inserted in the filter before the current block. The metadata of each slot consumes *r* bits, one bit for each *isOccupied* and *isRunEnd* metadata, and configurable *f*-bits and *t*-bits for the fixed counter and the slot-specific label respectively.

Moreover, the new counting scheme in MQF is simplified compared to that of the CQF. MQF defines the required memory for any item based solely on its count. Therefore, an accurate estimation of the required memory for any dataset can be done extremely quickly by an approximate estimation of data distribution(15)(16). This is unlike CQF which needs to add a safety margin to account for the special slots used by the counter encoding technique since it is impossible to estimate the number of these slots.

Regarding the speed of insertions and queries, MQF is slightly faster than CQF (Figure 3). This could be explained partially by the lower load factor of MQF and partially by the simplicity of the coding/decoding scheme of its counting system. Both MQF and CQF are faster than CMS unless the target FPR is really high (e.g. FPR > 0.1) (Figure 3). CMS controls its FPR by increasing the number of its hash tables requiring more time for insertions and queries to happen. In comparison, quotient filters use always one function but with more hash-bits to control the FPR, with a minimal effect on the insertion/query performance (Figure 3). With high FPR (e.g. FPR = 0.1), CMS uses fewer hash functions and is better performing than MQF. A quotient filter or CMS with a FPR = *δ* should have the same probability of item collisions. However, the quotient filter will be more accurate because CMS has another type of error with a probability (1-*δ*), which incorrectly increases the count of its items. This error is a “bounded error” with a threshold that inversely correlates with the width of the CMS(12). In another sense, some applications might deploy CMS with a smaller table’s width to be more memory efficient than MQF if the application can tolerate a high bounded error.

Buffered MQF can trade some of the speed of insertions and queries for significant memory reduction by storing data on disk. The buffered structure was developed to make use of the optimized sequential read and write on SSD. The buffered structure processes most of the insertion operations using the bufferMQF that resides on memory, thereby limiting the number of access requests to the MQF stored on the SSD hard drive. Sequential disk access happens when the bufferMQF needs to be merged to the disk. This approach is very efficient for insertions but not for random queries which require more frequent SSD data access. In k-mer analysis of huge raw datasets, buffered MQF can be used initially to filter out the low abundant k-mers (i.e. likely erroneous k-mers), then an in-memory MQF holding the filtered list of k-mers could be used for subsequent application requiring frequent random queries. This allows multistage analyses where a first pass eliminates likely errors (17–20).

CMS is commonly used for online or streaming applications as long as their high error rate can be tolerated (21). MQF has a better memory footprint in the approximate mode for lower error rates and thus can compute with CMS for online applications. A major advantage of quotient filters compared to CMS is the dynamic resizing ability in response to the growing input dataset (4). The buffered version of MQF can be very useful when the required memory is still bigger than the available RAM. We should, however, notice that online applications on MQF cannot make use of the memory optimization that could be achieved with an initial estimation of the filter parameters. A new version of the Khmer software that replaces CMS with MQF proves the new data structure more efficient in real-life applications. The MQF version is faster than the one with CMS unless the target FPR is high. Also, MQF is always more accurate than CMS although both structures have the same FPR. This behavior of CMS is due to the high error bound of its counts.

MQF comes with a novel labeling system that supports associating each k-mer with multiple values. There are two types of labels: Internal labels adjacent to each item to achieve the best cache locality but has a fixed size and thus practically useful when a small size label is needed. The second labeling system is to label the k-mers with one or more labels stored in external arrays while using the k-mer order in the MQF as an index. External labeling is very memory efficient mimicking the idea of the minimal perfect hash function (MPHF) (10,11). MPHF undoubtedly has the least memory requirement of all the associative data structures (11). However, MQF has better performance in both the construction and query phases. For construction, both structures require initial k-mer counting. MQF needs just an extra O(N) operation to update the block labels where N is the number of its unique k-mers. MPHF has to read then rehash the list of unique k-mers possibly more than once which makes it slower than MQF. For query operations, MQF is 10x faster regardless of the load factor of MQF (supplementary figure 2).

Furthermore, MQF offers more functionality and has fewer limitations than MPHF. MQF is capable of labeling a subset of its items which saves significant space for many applications. For example, k-mer analysis applications may want to only label the frequent k-mers, as an intermediate solution between pruning all the infrequent k-mers and labeling all the k-mers. Moreover, MQF allows online insertions and deletions of items as well as merging of multiple labeled MQFs (See the methods) while MPHF - which doesn’t store the items - needs to be rebuilt over the whole dataset, which requires reading and rehashing the datasets. Furthermore, MQF can be exact, while MPHF has false positives when queried with novel items that don’t belong to the indexed dataset.

## Conclusions

MQF is a new counting quotient filter with a simplified encoding scheme and an efficient labeling system. MQF adapts well to a wide range of k-mer datasets to be more memory and time-efficient than its predecessor in many situations. A buffered version of MQF has a fast insertion algorithm while storing most of the structure on external memory. MQF combines a fast access labeling system with MPHF-like associative functionality. MQF performance, features, and extensibility make it a good fit for many online algorithms of sequence analysis.

## Methods

### MQF Data structure

MQF has a similar structure to CQF with a different scheme of metadata that enables different counting and labeling systems (Figure 4). Like the CQF, the MQF requires 2 parameters, *r* and *q*, and creates an array of 2^*q*^ slots; each slot has *r*-bits. In Q MQF, *Q_i_* is the slot at position i where *i* = 1… 2^*q*^. The MQF maintains the block design of CQF where each block has 64 slots with their metadata and one extra byte of metadata called *Offset* to enhance the query of items(4). Both MQF and CQF have two metadata bits to accompany each slot: *isRunEnd_i_* and *isOccupied_i_*. In the MQF, each slot *i* has extra metadata, a fixed-size counter with a value (*F_i_*) and a configurable size (F_size_). There are also two optional fixed-size parts of metadata allocated to allow different styles of labeling. Every slot has specific labeling (*ST_i_*) with a configurable size (*ST_size_*>=0), and every block (*j*) has an optional space of a configurable size designed to store the number of items in the previous blocks.

The MQF uses the same insertion/query algorithm of CQF (4). In brief, suppose item *I*, repeated *c* times, is to be inserted into Q. A hash function *H* is applied to *I* to generate a *p*-bit fingerprint (*H*(*I*)). *H*(*I*) value is split into two parts, a quotient and remainder. The quotient (*q_i_*) is the most significant *q* bits while the remainder (*r_i_*) is the remaining least significant r bits. The filters store *r_i_* in a slot *Q_j_* where *j*≥*q_i_*. One or more slots can be used to store the count of the same item. If the required slots for the item or its count are not free, all the consecutive occupied slots starting from this position will be shifted to free the required space. All items having the same *q* are stored into consecutive slots and are called a run. Items in the run are sorted by *r_i_*, and *isRunEnd* of the last slot in the run is set to one. *isOccupied*(*q_i_*) is set to one if and only if there is a run for *q_i_*. Therefore, there is one bit set to one in each *isOccupied* and *isRunEnd* for each run. To query item *I*, a Rank and Select method is applied on the metadata arrays to get the run start and end for *q_i_*. Then all the items in the run are searched linearly for the slot containing *r_i_*. The subsequent one or more slots can be decoded to get the count of item *I*. CQF uses a special encoding scheme to recognize these counting slots but MQF utilizes the fixed-counter metadata element (see below).

### Counting scheme

MQF uses two types of counters for storing the values of the count (*c*): A small fixed-size space (*F_i_*) is slot specific and used to store the count of the item in its own slot if this count is smaller than *F_max_*, where *F_max_* is the maximum possible value for the fixed space. A variable size space (*V_i_*) is composed of one or more slots next to the item’s slot and is used to store larger values. For an item with high count *c*, the number of required slots for *V_i_* is calculated as 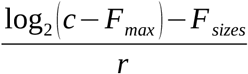 slots. The *F_i_* spaces of this item’s slot and its *V_i_* slots are used to mark the last slot for the item where all of them will be saturated to *F_max_* except the last one (Figure 5). This counting scheme can be summarized into 2 rules:

*Rule 1:* MQF requires *F_i_* < *F_max_* if and only if *i* is the index of the item’s last slot.
*Rule 2:* If *c* < *F_max_*, *c* is stored in *F_i_* only. Otherwise, *c* - *F_max_* is stored in *V_i_*

**Figure 5:**
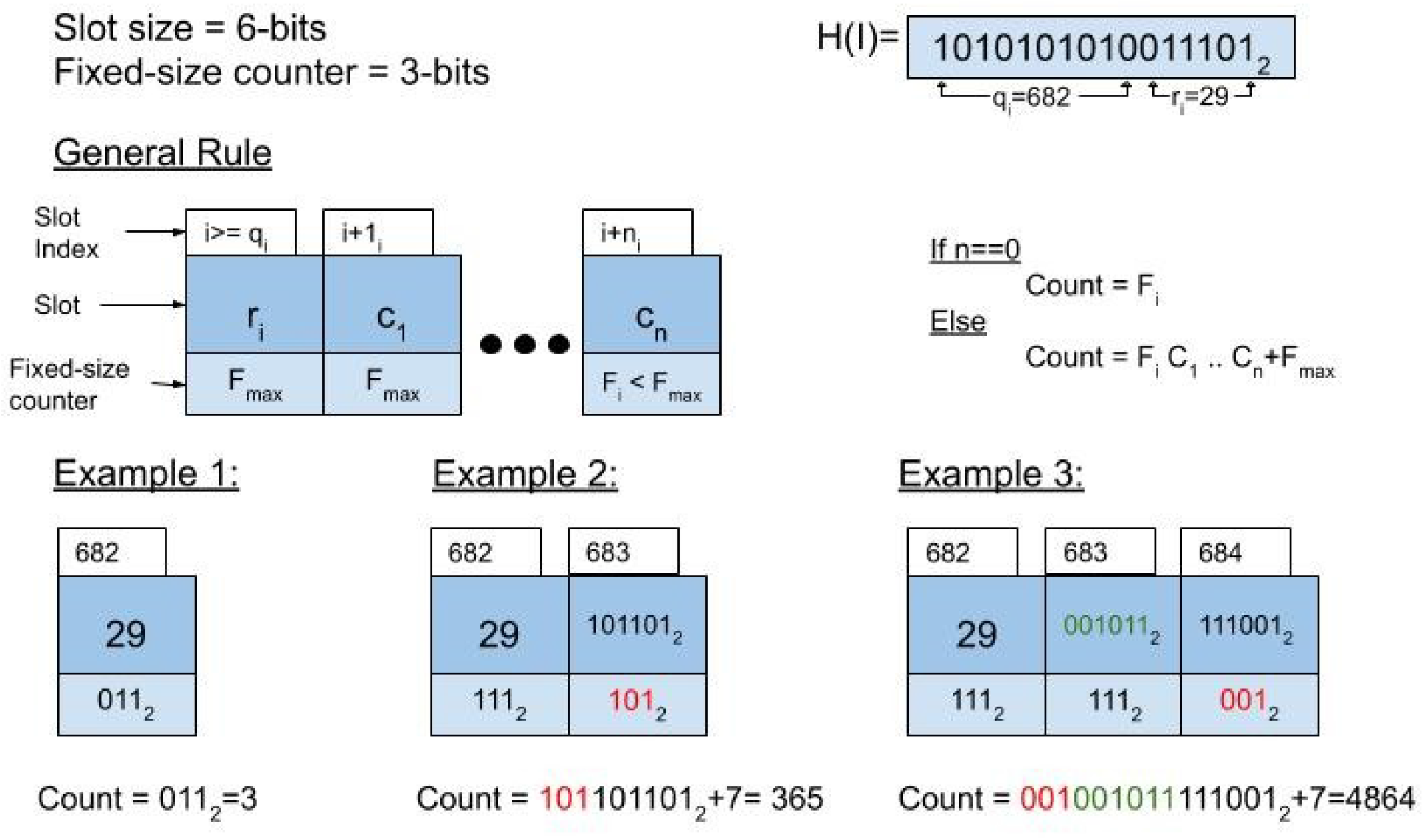
MQF counters encoding scheme. Items and their counts are stored in n slots and n fixed counter as shown in the general rule. Each example stores the same item but different count (count = 3, 365, or 4864).

In comparison to the CQF, the MQF does not use special slots to resolve ambiguities, which is more memory efficient. The counter encoding algorithm is described in Supplementary Figure 3.

## Parameter Estimation

For offline counting applications, the MQF parameters (q, r, F_size_) can be even more optimized for each dataset to create the most memory-efficient filter that has enough slots to fit all unique items and their counts. The *q* parameter defines the number of slots (N) in MQF where *q*=log_2_ (*N*). The required numbers of slots for items and their count can be estimated from the cardinality of the target dataset, as with CQF. The *r* parameter is calculated from the equation *r* = *p-q* where *p* is the total number of hash-bits used to represent each item. In the exact mode, *p* equals the exact output of a reversible hash function. In the inexact mode, *p* is controlled by the target FPR(*δ*) according to the equation 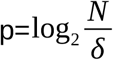 described before (4). The F_size_ parameter defines the size of the fixed-size counter. This is critical because if a given MQF has too few slots for items in a dataset, the bigger MQF would have to double the number of slots causing a big jump in the memory requirement. To avoid that jump, MQF can use larger fixed size counters to decrease the number of slots required in counting on the expense of a slight increase in the slot size.

## Labeling System

MQF can map each item to its count as well as other values, which we call “labels”. Labels in MQF have two different systems. An internal labeling system stores the associated value for every key in the data structure, like a hash table. This label has a fixed size defined at the initialization of the MQF and is practically useful when a small size label is needed (e.g. one or two bits). The second labeling system labels the block. We use this label to store the number of items inserted in the MQF before each block. This enables labeling the items of the filter by separate arrays matching the order of the items in the filter, a behavior that can act as a minimal perfect hash function (11). The naive way to compute the items’ order is to find the item in the MQF and iterate backward until the beginning of the filter to count the number of the preceding items, which is an O(N) operation. The MQF stores the number of items that exist before each block; therefore, the MQF iterates only to the beginning of each block, which is an O(1) operation. The number of previous items for each block is computed after the MQF is constructed. Any additional insertions or deletions of items would only require re-calculation of the block label values with no need to re-analyze the original data. Moreover, labeled MQFs can be updated by merging multiple labeled MQFs and their external labeling arrays. External label arrays need to be merged after merging the labeled MQFs. To do so, the new items’ order is recomputed in the final MQF. Then, labels in the input external arrays can be copied into a new external array according to the new item order. Such a function has to consider resolving the conflicts of items happening in multiple-input MQF and labeled by different external labels (Figure 6).

**Figure 6:**
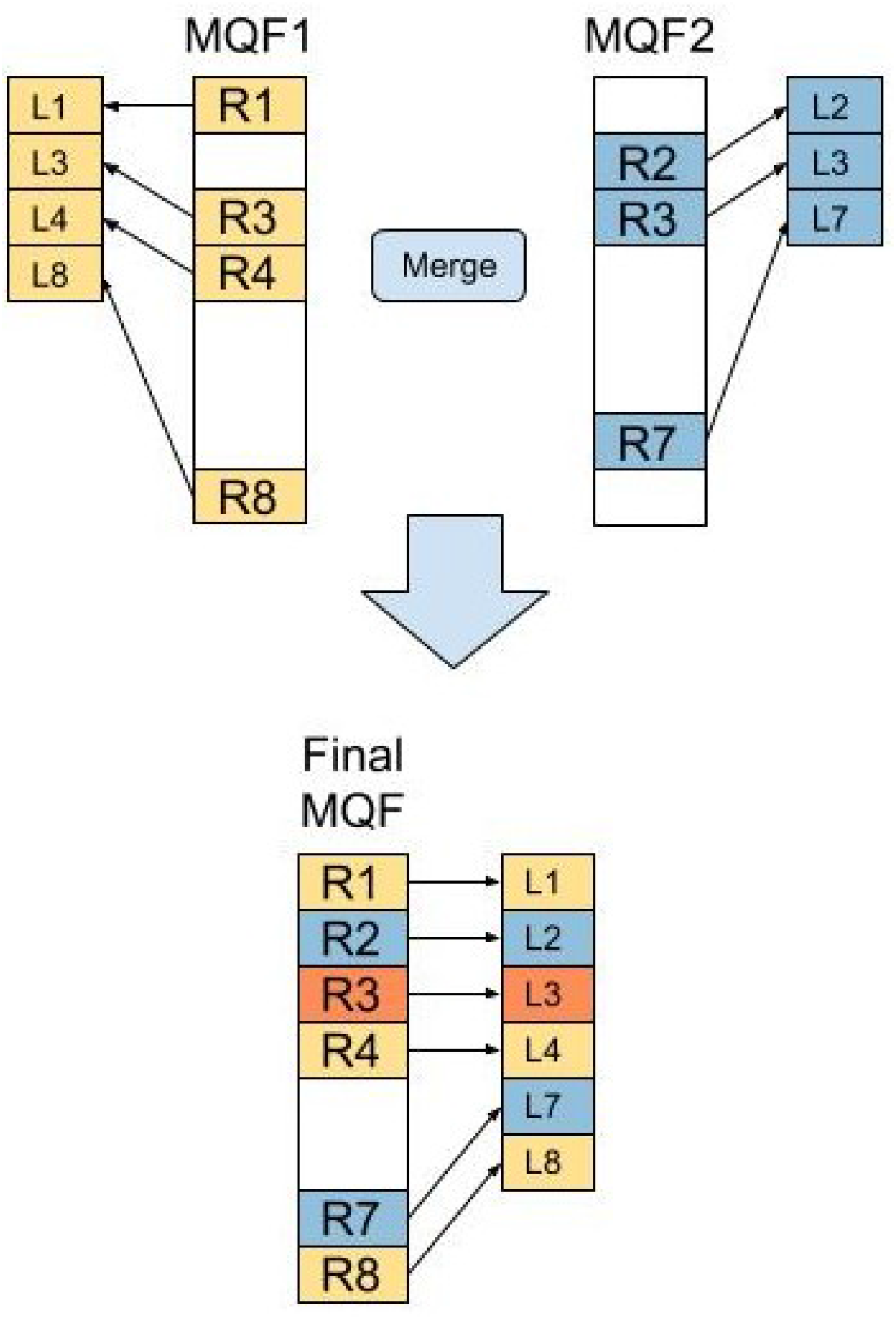
Merging MQFs with external labels. R_i_ is the remaining part of item i, and Ti is the external label of the item. Merging the input MQF produces a final MQF with a new order of its member items. All labels in the input external arrays are copied into a new external array according to this new order of the items. However, the implementation of the merge function has to resolve the conflict of R3 labels which exist in both input structures with two labels.

## Buffered MQF

The Buffered MQF is composed of two MQF structures: a big structure stored on SSD called onDiskMQF, and an insertion buffer stored in the main memory called bufferMQF. OnDiskMQF uses stxxl vectors(22) because of the performance of their asynchronous IO. The bufferMQF is used to limit the number of accesses on the OnDiskMQF and change the access pattern to the on-disk structure from random to sequential. As shown in the insertion algorithm in Figure 7, all the insertions are done first on bufferMQF; when it is full, the items are copied from bufferMQF to OnDiskMQF, and bufferMQF is cleared. The copy operation edits the onDiskMQF in a serial pattern which is preferred while working on SSD because many edits will be grouped together in one read/write operation. Figure 8 shows the query algorithm. The queried items are inserted first to temporary MQF and sequential access is done to query the items from the OnDiskMQF. The final count is the sum of the bufferMQF and the ondiskMQF.

**Figure 7:**
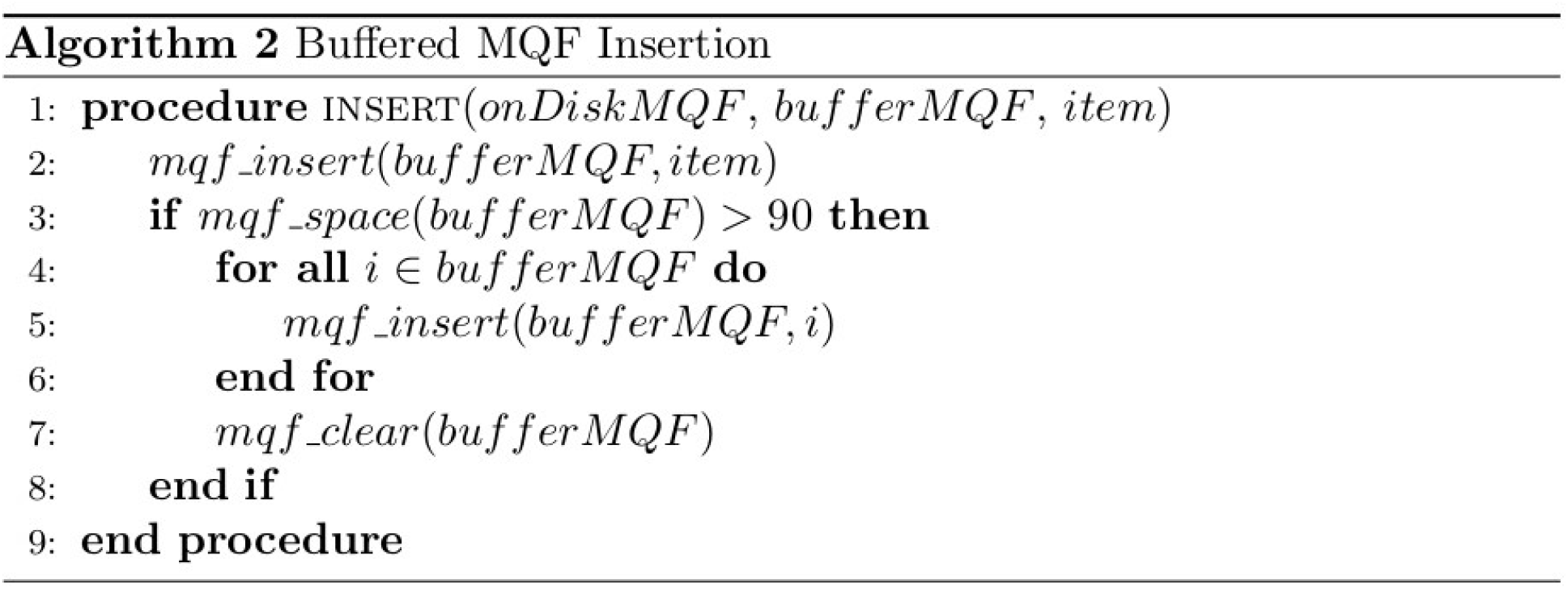
Buffered MQF insertion algorithm. Insertion Algorithm for inserting items in the Buffered MQF. It inserts the item in the in-memory data structure. The on-memory structure is merged into the on-disk structure when it is filled.

**Figure 8:**
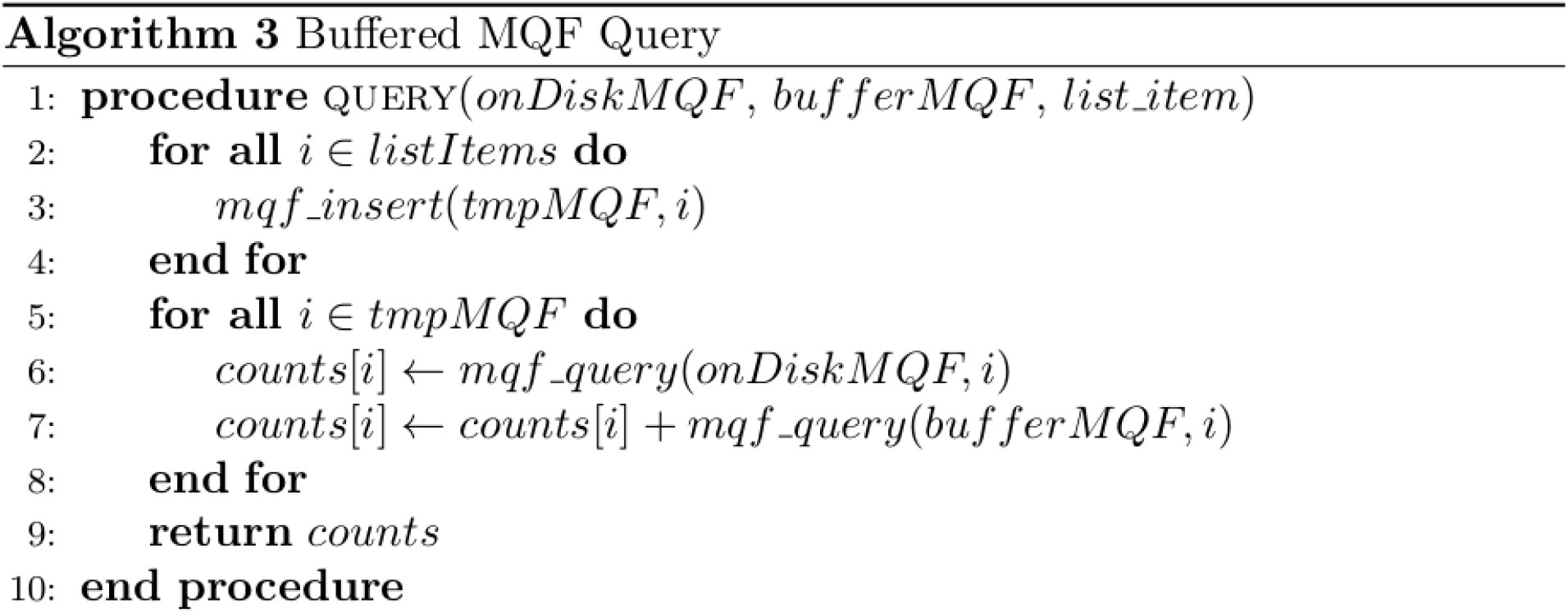
Buffered MQF query algorithm. Query algorithm for retrieving counts for a list of items in the Buffered MQF. First, insert all the items in the list into a temporary MQF. Second, iterate over the list of items in the temporary MQF and query both the in-memory and on-disk structures.

## Experimental Setup of Benchmarking

Five datasets were used in the experiments to cover most of the bioinformatics applications. Three datasets called z2, z3, and z5 were simulated to follow Zipfian distribution using three different coefficients: 2, 3, and 5 respectively. The bigger the coefficient the more singletons in the dataset (23). A fourth dataset was simulated from a uniform distribution with a frequency equal to 10. One more dataset, named k-mers, represented real k-mers generated in the ERR1050075 RNA-seq experiment from humans(24). Experiments were conducted to compare the performance, memory, and accuracy of MQF with the state-of-the-art counting structures CQF, CMS, and MPHF. Unless stated otherwise, CQF and MQF used the same number of slots, and the same slot size while the fixed counter of MQF was set to two. The slot size was calculated to achieve the target FPR as described in the parameter estimation section (see Methods). To create comparable CMS, the number of the tables in the sketches was calculated using ln 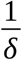 as described before (12). The table width was calculated by dividing the MQF size by the number of tables. The MPHF was created using the default options in the BBhash repo (https://github.com/rizkg/BBHash). An Amazon AWS t3.large machine with Ubuntu Server 18.04 was used to run all the experiments. The instance had 2 VCPUS and 8GB RAM with a 100GB provisioned IOPS SSD attached for storage. All codes used in the experiments can be accessed through the MQF GitHub repository (https://github.com/dib-lab/2020-paper-mqf-benchmarks).

## List of abbreviations

MQF: mixed-counters quotient filter.
CQF: counting quotient filter.
FPR: false positive rate.
CMS: count-min sketch.
MPHF: Minimal Perfect Hash Functions

## Declarations

### Ethics approval and consent to participate

Not applicable

### Consent for publication

Not applicable

### Availability of data and materials

The datasets used in Benchmarking are available in the “2020-paper-mqf-benchmarks” repository. https://github.com/dib-lab/2020-paper-mqf-benchmarks

### Authors’ contributions

TAM and MS developed theoretical formalism. MS carried out the implementation and benchmarking. TAM conceived the original idea and supervised this work. All authors contributed to the writing of the manuscript.

## Acknowledgements

Not applicable

