## Supplementary Figures for "MQF and buffered MQF: Quotient filters for efficient storage of k-mers with their counts and metadata"

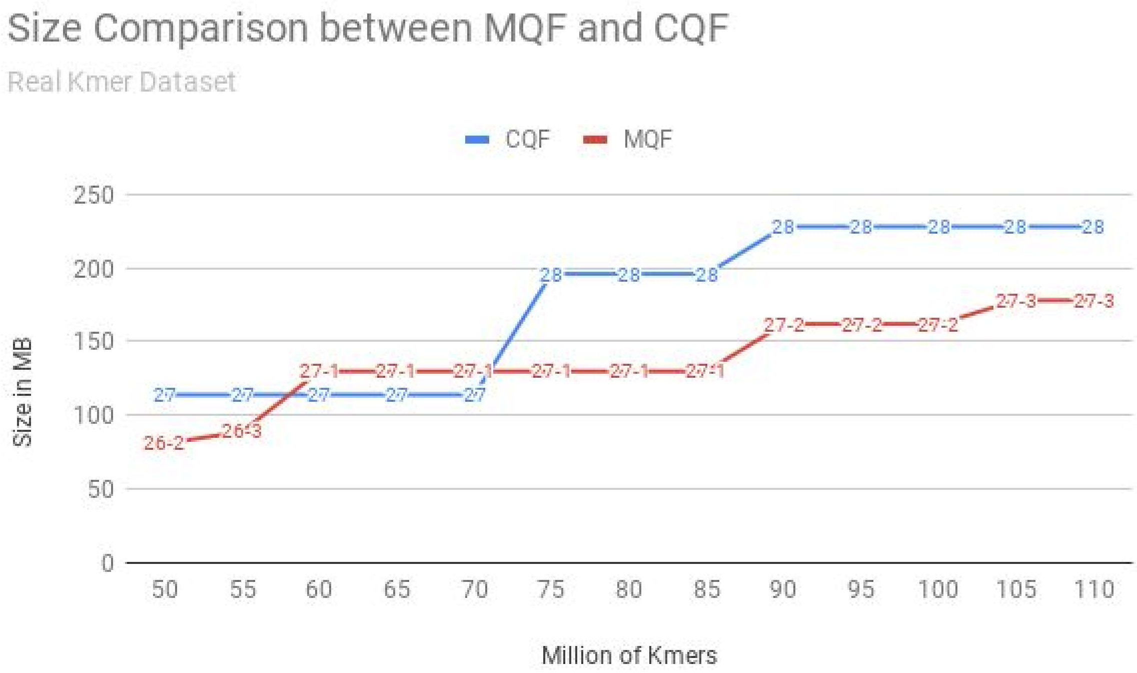


**Supplementary Figure 1: Detailed Memory Consumption Comparison.** The numbers on the CQF curve are the used Q sizes. While the numbers on MQF are (Q size-fixed counter size).

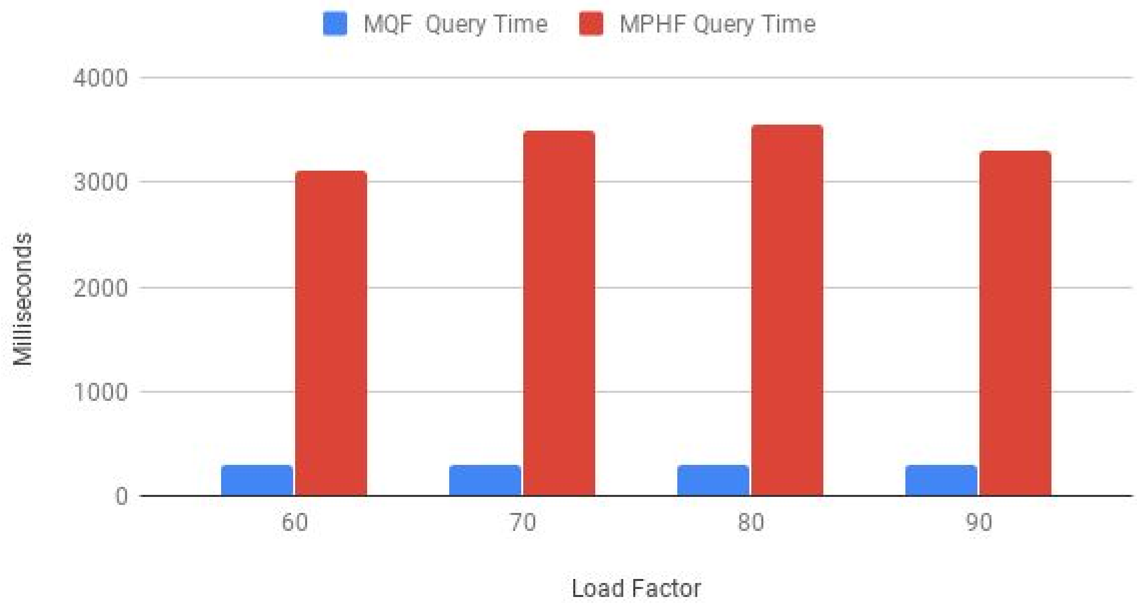


**Supplementary Figure 2: Query performance Comparison of MQF and MPHF**. The query times of 35M existed kmers were measured for MQF structures with 60%, 70%, 80%, and 90% load factors and MPHF structures storing matching datasets.

**Algorithm 1** Counters Encoder

**procedure** ENCODE($Q$, $r_i$, $count$ ,$start$)
  $base \leftarrow 2^{Q.r}$ ▷ Q.r is #bits in the slot
  $fcountMax \leftarrow 2^{Q.f} - 1$ ▷ Q.f is #bits in the fixed-size counter
  $stack \leftarrow \phi$
  **while** $count > fcountMax - 1$ **do**
    $stack.push(count\%base)$
    $count = count >> Q.r$ ▷ bit shift operation
  **end while**
  $stack.push(r_i)$
  $i \leftarrow start$
  **while** $stack \neq \phi$ **do**
    $Q_i.r \leftarrow stack.pop()$ ▷ Slot of index i
    $Q_i.f \leftarrow fcountMax$ ▷ fixed-size counter of index i
    $i \leftarrow i + 1$
  **end while**
  $Q_i.f \leftarrow count$
**end procedure**

**Supplementary Figure 3: Counters encoder algorithm.** The Algorithm encodes the item and its count into one or more slots. The first slot is reserved for the item's remaining, and the variable number of slots follows to encode the item's count.

| Distribution | CQF | MQF |
|---|---|---|
| Z2 | 402 | 671 |
| Z3 | 161 | 161 |
| Z5 | 134 | 134 |
| Kmers | 402 | 671 |
| Uniform | 402 | 1275 |

Supplementary Table 1: Number of items (in millions) inserted to achieve a 90% load factor in compact hash tables.
